# PD-1 suppresses the maintenance of cell couples between cytotoxic T cells and tumor target cells within the tumor

**DOI:** 10.1101/443788

**Authors:** Rachel Ambler, Grace L. Edmunds, Giulia Toti, David J. Morgan, Christoph Wülfing

## Abstract

CD8^+^ T cell killing of tumor cells is suppressed by the tumor microenvironment. Inhibitory receptors, prominently PD-1, are key mediators of this suppression. To discover cellular defects triggered by tumor exposure and associated PD-1 signaling, we have established an ex vivo imaging approach to investigate CD8^+^ tumor infiltrating lymphocytes (TILs) interacting with tumor targets. Whilst TIL:tumor cell couples formed effectively, couple stability deteriorated within 1-2 minutes. This was associated with excessive cofilin recruitment to the cellular interface, coincident deterioration of f-actin structures, increased TIL locomotion, and impaired tumor cell killing. Diminished engagement of PD-1 within the tumor, but not acute ex vivo blockade, partially restored cell couple maintenance and killing. PD-1 thus suppresses TIL function by inducing a polarization-impaired state.

## Introduction

Cancer cells are commonly recognized by the immune system and immune cells constitute a large part of the tumor mass. Cytotoxic T lymphocytes (CTL) are capable of directly killing cancer cells. However, their killing ability is widely suppressed after they have entered the tumor, where they are known as tumor infiltrating lymphocytes (TIL). A number of key contributors to in vivo tumor-mediated immune suppression have been characterized. Most prominently, tumor-reactive T cells increase expression of inhibitory receptors, including CTLA-4, PD-1, LAG3, TIGIT and TIM3 (1, 2). Monoclonal antibody blockade of CLTA-4 and PD-1 has yielded substantial clinical success in enhancing the anti-tumor immune response (3). Additional mechanisms of tumor suppression include recruitment of tolerogenic immune cells, notably regulatory CD4^+^ T cells (Tregs) into the tumor (4, 5) and the expression of suppressive soluble mediators, such as adenosine and prostaglandin-E_2_ (PGE_2_)(6-8). While it is clear that tumor cell killing is diminished due to the immunosuppressive tumor microenvironment it is still uncertain as to which cellular steps in target cell killing are impaired in TILs and how such impairment is controlled by the established mediators of tumor-mediated immune suppression.

PD-1 is a critical mediator of tumor-mediated immune suppression and is upregulated in response to continuous antigen exposure. In persistent viral infections PD-1 signaling maintains an exhausted phenotype among CD8^+^ T cells (9). PD-1 engagement has been shown to have a wide variety of effects on T cell activation. These include activation of the phosphatase PTEN (10), recruitment of the phosphatase SHP-2 (11, 12), suppression of sustained activation of Akt and Ras pathways with consequences for cell cycle regulation (13), inhibition of glycolysis along with promotion of lipolysis and oxidative metabolism (14) and upregulation of the proapoptotic protein Bim (15). The T cell stop signal upon antigen presenting cell contact (16) and the formation of stable cell couples (17) were inhibited. However, it remains unresolved whether and how PD-1 signalling disrupts the cellular events required for cytolytic killing.

The killing of target cells by CTLs requires carefully orchestrated steps of cellular reorganization (‘polarization’). CTLs must bind to the target cells, use actin polymerization to stabilize the cellular interface, relocate the MTOC from behind the nucleus to the center of the cellular interface, and finally release the contents of cytolytic granules directed towards the target cell (18, 19). To investigate how such polarization may be altered when TILs interact with their tumor target cells, we have adapted a well-established model of antigen-driven tumor recognition to ex vivo live cell imaging approaches.

A renal carcinoma cell line (Renca) expressing the influenza virus haemagglutinin (HA) as a neo-antigen (RencaHA) is recognized by CD8^+^ T cells from Clone 4 TCR transgenic mice (20). When adoptively transferred into a RencaHA-tumor bearing BALB/c mouse naïve Clone 4 T cells are primed within the tumor-draining lymph nodes, infiltrate the tumor and acquire a suppressed phenotype that is characterized by diminished cytokine secretion, killing ability and enhanced inhibitory receptor expression (21). Here we have adoptively transferred in vitro activated Clone 4 CD8^+^ T cells, which had been retrovirally transduced to express fluorescently tagged signaling proteins, into RencaHA tumor-bearing mice. These transduced Clone 4 T cells homed to the tumor and acquired a suppressed phenotype. They were reisolated for an ex vivo investigation of their interaction with RencaHA tumor cells. Clone 4 TILs displayed impaired cell couple maintenance with their tumor targets that was driven by diminished F-actin stability caused by excessive cofilin recruitment to the TIL target cell interface, resulting in impaired tumor cell killing. PD-1 expression on Clone 4 TILs was elevated. Blocking PD-1 engagement in vivo in the tumor microenvironment, but not acutely in vitro, enhanced tumor clearance and improved the maintenance of TIL cells. Thus we have we have discovered a cellular mechanism of tumor-mediated immune suppression: Impaired maintenance of T cell polarization diminished TIL killing ability and was regulated by the inhibitory receptor PD-1.

## Results

### The cytolytic ability of CD8^+^ T cells is diminished by tumor infiltration

To investigate the killing of tumor target cells by CD8^+^ TILs at subcellular resolution, we adapted a mouse model of neo-antigen CD8^+^ T cell tumor recognition to enable ex vivo live cell imaging approaches (Fig. 1A). RencaHA renal carcinoma cells express the haemagglutinin (HA) protein from influenza A/PR/8/H1N1 as a neo-antigen. The Clone 4 TCR recognizes the dominant H2-K^d^ – restricted HA peptide 518-526 (IYSTVASSL). In all experiments, CD8^+^ T cells from Clone 4 TCR transgenic mice were activated in vitro with HA peptide-pulsed splenocytes, retrovirally transduced to express GFP-tagged proteins of interest, and FACS sorted for minimal sensor expression (2μM) which is close to endogenous protein levels (22). To generate Clone 4 TILs, 4×10^6^ Clone 4 CD8^+^ T cells were adoptively transferred into BALB/c mice bearing subcutaneous RencaHA tumors of approximately 500mm^3^. After 96h the GFP-positive TILs were isolated from the tumors by FACS sorting. We used two cell populations as fully active control cytolytic effectors: Clone 4 T cells were either isolated from the spleen of tumor-bearing mice or directly activated in vitro with HA peptide-pulsed splenocytes (‘CTL’) as no apparent cell biological differences between these two control populations were observed (see below).

**Fig. 1.**
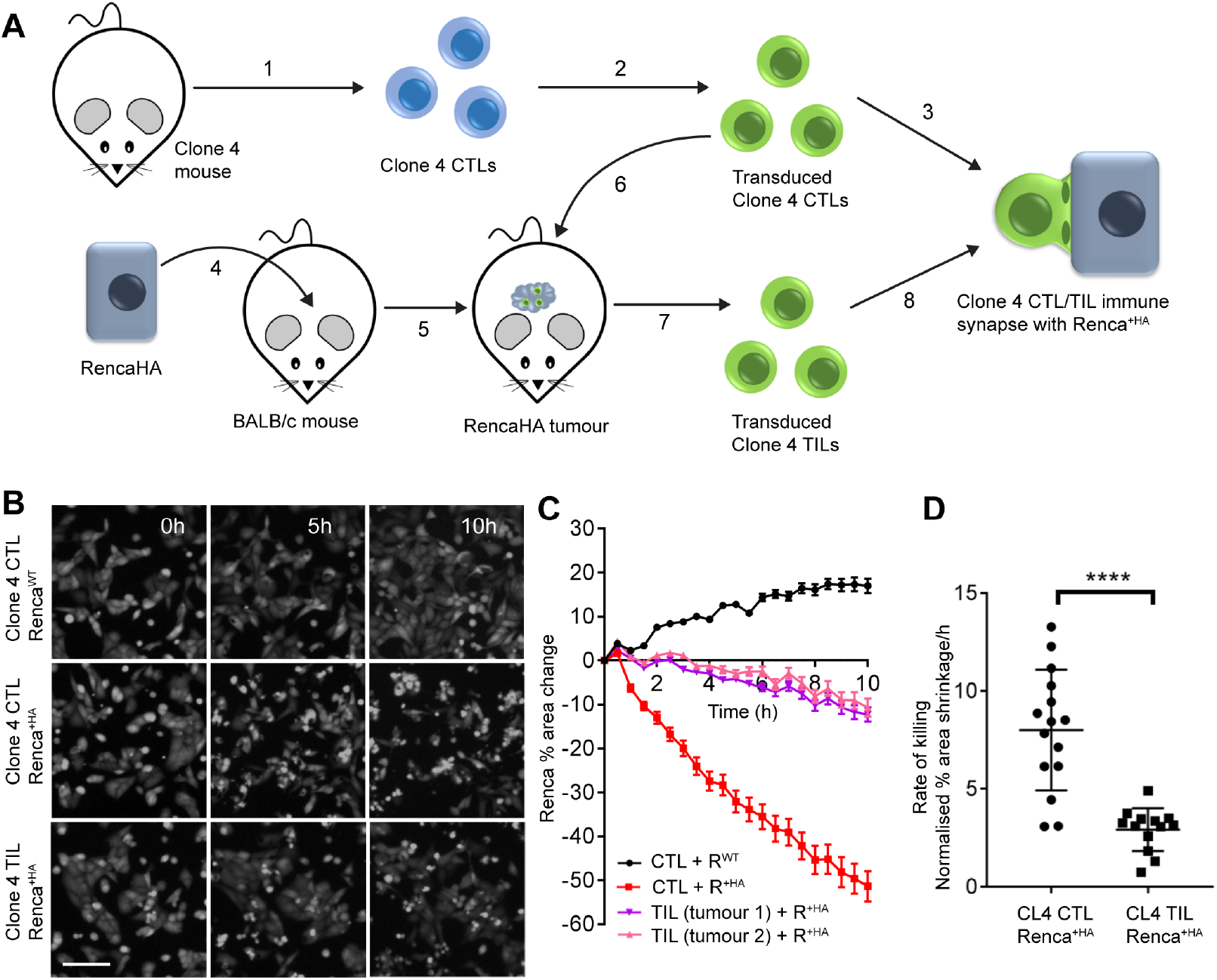
Tumour exposure impairs CD8^+^ T cell killing ability. (A) Schematic of the experimental design enabling comparative imaging of signalling events in Clone 4 CTLs and TILs. 1 in vitro priming of Clone 4 T cells with APC and peptide, 2 retroviral transduction of Clone 4 CTLs for the expression of fluorescent proteins, 3 imaging of the interaction of FACS-sorted fluorescent Clone 4 CTLs with Renca^+HA^ cells, 4 inoculation of BALB/c mice with RencaHA cells, 5 establishment of subcutaneous tumor, 6 adoptive transfer of FACS-sorted fluorescent Clone 4 CTLs, 7 isolation of fluorescent Clone 4 TILs from the tumor, 8 imaging of the interaction of fluorescent Clone 4 TILs with Renca^+HA^ cells (B) Representative images over time showing Clone 4 CTL and TIL killing of CTV labelled Renca cells pulsed with (Renca^+HA^) or without (Renca^WT^) HA antigen. Scale bar shows 100μm. (C) Percentage change in CTV labelled Renca^+HA^ and Renca^WT^ cell area over time following incubation with Clone 4 CTL and TIL, from representative killing assay shown in (B). Mean ± SEM, minimum of 3 fields of view per time point. (D) Average rate of CTV labelled, HA peptide pulsed Renca^+HA^ death following incubation with Clone 4 CTLs and TILs, calculated as percentage area decrease per hour. Each point is a separate CTL or TIL preparation, gathered over seven independent experiments. Mean ± SD, **** p < 0.0001, Student’s t test.

To confirm that RencaHA tumors suppress the cytolytic capability of Clone 4 CTLs, we developed an imaging-based ex vivo killing assay. Renca cells plated in a glass-bottom imaging plate were pulsed with 2 μg/ml HA peptide (Renca^+HA^) and were then overlayed with Clone 4 CTLs or TILs. Over a ten-hour period CTLs mediated efficient target cell lysis, decreasing the imaging area covered by Renca^+HA^ cells by 50% at a rate of 8±3 %/h (Fig. 1B to D). Non-HA peptide-pulsed Renca cells continued to proliferate which confirmed cognate antigen dependence. In contrast, Clone 4 TILs displayed a substantial loss of cytolytic ability. The imaging area covered by Renca^+HA^ cells was reduced by only 11% after 10 hours, at a significantly (p<0.0001) reduced rate of 3±1 %/h (Fig. 1B to D). To corroborate antigen-specific target cell killing by Clone 4 CTLs in vivo, BALB/c splenocytes were labeled with high and low amounts of CellTrace Violet, pulsed with or without HA peptide respectively, and injected intravenously into BALB/c mice which had previously received 4×10^6^ Clone 4 CTLs. After 12 hours HA pulsed splenocytes were specifically lost (fig. S1).

### TILs are deficient in MTOC polarization and cell couple maintenance

In cytotoxic T cells relocation of the MTOC from behind the nucleus to the target cell interface upon cell coupling is critical for lytic granule release and cytolytic activity. To visualize Clone 4 CD8^+^ T cell MTOC polarization toward the Renca target cell interface, Clone 4 CTLs and TILs expressing tubulin-GFP were imaged during their interaction with Renca^+HA^ cells in three dimensions over time by spinning disk confocal microscopy. Whereas 100% of CTLs relocated the MTOC to the cellular interface within 7min, there was a significant (p<0.01) decrease in MTOC relocation among TILs (67±8%)(Fig. 2A to C, fig. S2).

**Fig. 2.**
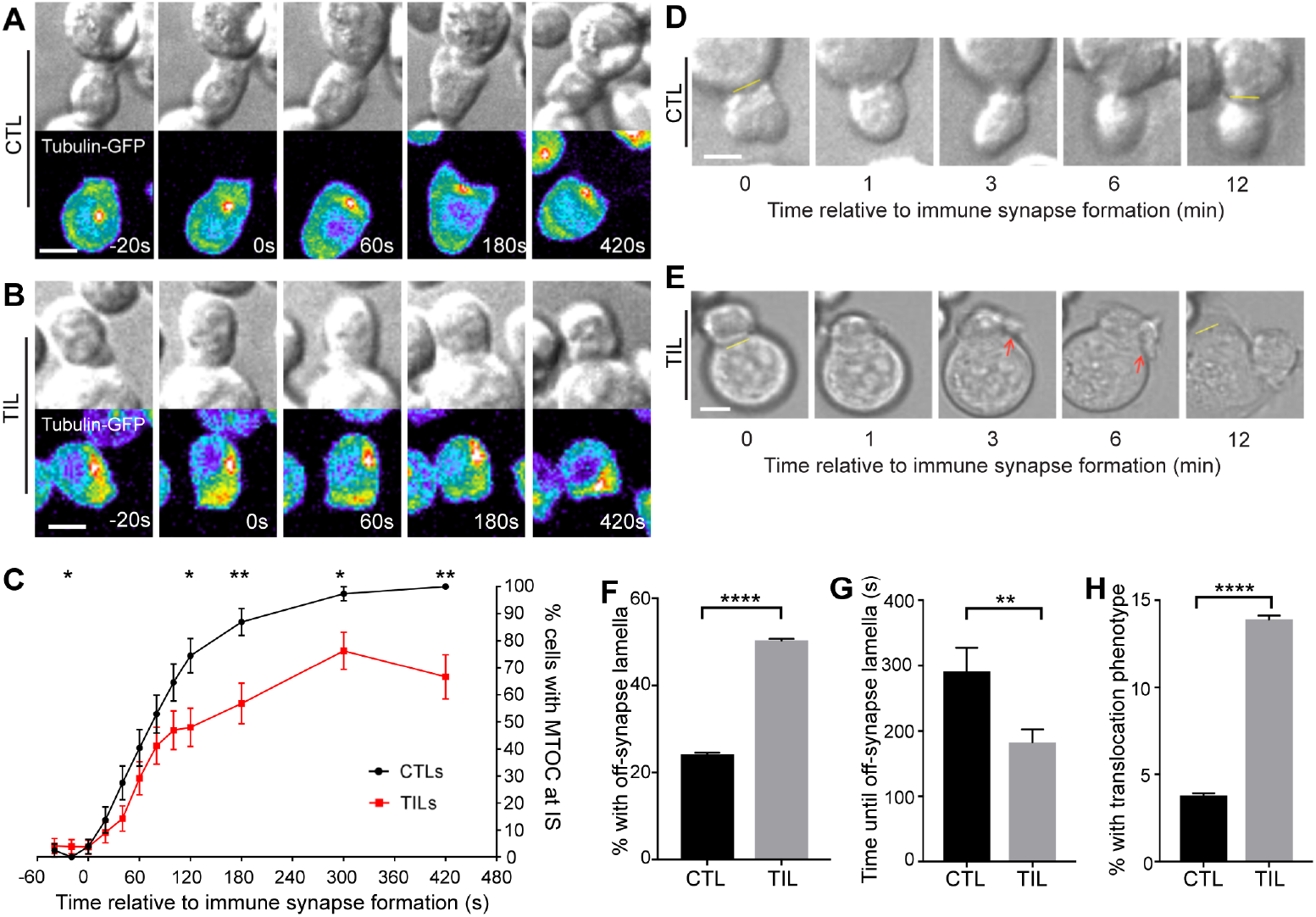
Diminished polarisation and cell couple stability of Clone 4 TILs. (A and B) Representative interaction over time of tubulin-GFP transduced Clone 4 CTLs (A) and TILs (B) with HA peptide pulsed Renca^+HA^ cells. Shown are matching DIC and maximum GFP projection panels with a false colour GFP intensity scale (blue to red). Scale bar shows 5μm. Complete interactions in Videos S1, S2. (C) Percentage over time of tubulin-GFP transduced Clone 4 CTLs and TILs, as in panels A/B, displaying contact between the MTOC and immune synapse during interaction with HA pulsed Renca^+HA^ cells. Tight cell coupling: time 0s. Mean ± SEM from 51 CTLs and 55 TILs from ≥2 separate experiments. (D and E) Representative interaction over time of CTLs (D) and TILs (E) with HA peptide pulsed Renca^+HA^ cells. Yellow line indicates original position of tight-cell couple formation on Renca^+HA^ cell. Red arrows indicate direction of membrane protrusions, termed off-synapse lamellae. (F-H) Percentage of Clone 4 CTLs and TILs coupled to HA peptide pulsed Renca^+HA^ cells with off-synapse lamellae (F), time of first off-synapse lamella (G) and translocation of more than one immune synapse diameter around the Renca^+HA^ cell circumference (H). All data are mean ± SEM. 132 CTL from 8 separate experiments and 137 TIL from 4 separate experiments were analysed. * p < 0.05, ** p < 0.01, **** p < 0.0001; p values calculated using proportions z-test (C) or Student’s t test (F-H).

As another readout of cellular polarization we investigated Clone 4 CD8^+^ T cell morphology. Clone 4 CTLs formed symmetrical, stable interfaces with Renca^+HA^ target cells throughout 12 minutes of observation (Fig. 2D). Initial immune synapses between Clone 4 TILs and Renca^+HA^ target cells were also highly symmetrical, but polarization was diminished starting two minutes after tight cell coupling (Fig. 2E). To quantify this difference in Clone 4 CD8^+^ T cell polarization, we determined the occurrence of lamellae pointing away from the cellular interface (red arrow Fig. 2E). T cells continuously extend large lamellae, but whilst directing such lamellae towards the interface is expected to be stabilizing, directing them away from the interface (‘off-interface lamellae’) may weaken it. Off-synapse lamellae occurred in a significantly (p=<0.0001) higher proportion of TILs than CTLs (50±4% versus 24±0.5%, p<n)(Fig. 2F) and were observed earlier during cell coupling (183±20s versus 291±36%, p<n0.01)(Fig. 2G). Furthermore, TILs remained at the initial site of target cell binding less efficiently: 14±0.2% of TILs translocated across the target cell surface by at least one interface diameter from the initial site of target cell binding, a significantly (p<0.0001) higher percentage than observed in CTLs (4±0.4%)(Fig. 2H). Such T cell translocation across a tumor target cell surface has previously been described in the interaction of CTLs with explanted tumors (23, 24). While Clone 4 TILs could effectively establish cell couples with Renca target cells, they did not maintain cell couples well, moving away from the initial site of target cell binding in a substantial fraction of the population within a few minutes.

### Peripheral F-actin ring formation is destabilized in TILs

The immunological synapse is stabilized by an F-actin ring around its periphery. To determine whether impaired F-actin ring formation could underpin impaired maintenance of TIL cell couples, Clone 4 CTLs and TILs expressing the F-actin sensor F-tractin-GFP were imaged (Fig. 3, fig. S3). Within the first 20 seconds of tight cell coupling, 95±3% of CTLs and 80±6% of TILs successfully formed a peripheral F-actin ring (Fig. 3A, B, D, and E, fig. S3D and F) which is consistent with equally efficient cell coupling by TILs and CTLs. However, fewer TILs maintained the peripheral actin ring. At 180s and 420s, only 48±7% and 26±8% of TILs maintained a peripheral F-actin, compared with 88±5% and 61±8% of CTLs (p<0.0001, p<0.01, respectively)(Fig. 3D and E, fig. S3D and F). Tumor exposure did not significantly alter the amount of F-tractin-GFP recruited to the immune synapse (fig. S3H). Corroborating that in vitro generated Clone 4 CTLs are comparable to in vivo cytolytic Clone 4 T cells, peripheral F-actin ring formation was the same in Clone 4 CTLs and adoptively transferred Clone 4 CTLs re-isolated from the spleen of tumor bearing mice (fig. S3A to C). Intriguingly, the loss of Clone 4 TIL peripheral F-actin coincided with the average time of occurrence of off-interface lamellae (Fig. 2G) and the time at which MTOC polarization began to fail (Fig. 2C). Taken together, these data establish that tumor exposure induces a state of defective TIL polarization where couple maintenance becomes impaired two to three minutes after tight cell couple formation.

**Fig. 3.**
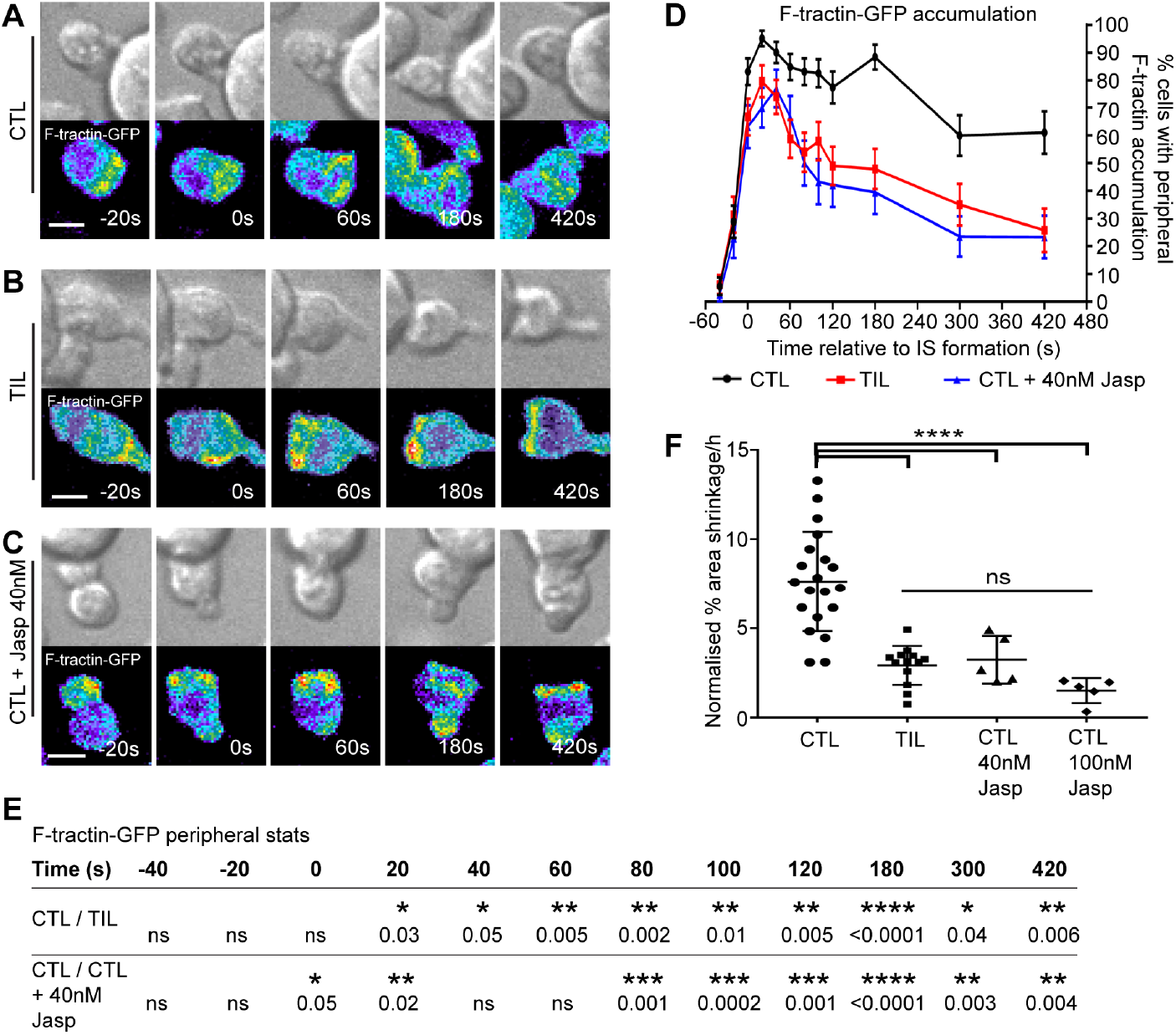
Impaired formation of the F-actin peripheral ring is associated with poor TIL killing ability. (A-C) Representative interaction over time with HA peptide pulsed Renca^+HA^ cells, of F-tractin-GFP transduced Clone 4 CTLs (A), TILs (B) and CTLs incubated with 40nM jasplakinolide for 30 minutes prior to and during cell coupling (C). Shown are matching DIC and maximum GFP projection panels with a false colour GFP intensity scale (blue to red). Scale bar shows 5μm. Complete interactions in Videos S3-S5. (D, E) The percentage over time of F-tractin-GFP transduced Clone 4 cells displaying peripheral F-tractin-GFP interface accumulation during interaction with HA peptide pulsed Renca^+HA^ target cells. Shown is data for CTLs, TILs, and CTLs treated with 40nM jasplakinolide for 30 minutes prior to and during coupling. Tight cell coupling: time 0s. Mean ± SEM, 60 CTLs, 51 TILs and 40 jasplakinolide treated CTLs, from ≥3 separate experiments. P values calculated by proportions z test (E). (F) Average rate of CTV labelled, HA peptide pulsed Renca^+HA^ cell death following incubation with Clone 4 CTLs, TILs and CTLs treated with either 40nM or 100nM Jasplakinolide 30 minutes prior to and during cell coupling. Data for CTLs and TILs previously shown in Fig. 1D. Mean ± SD, p values calculated by Student’s t test. * p < 0.05, ** p < 0.01, *** p < 0.001, **** p < 0.0001.

To investigate whether impaired cell couple maintenance results in loss of cytolytic effector function, Clone 4 CTLs were treated with a low concentration (40nM) of Jasplakinolide. At this concentration cell coupling was not impaired (25, 26), but the percentage of CTLs displaying a peripheral F-actin ring was markedly reduced to levels similar to those achieved among Clone 4 TILs (Fig. 3C to E, fig. S3F). Treatment with 40nM Jasplakinolide significantly (p=0.0001) reduced the rate of target cell lysis by CTLs to 3±1%/h to levels displayed by TILs (Fig. 3F). Treatment with a higher concentration (100nM) of Jasplakinolide impaired cell coupling (25) and further reduced killing (Fig. 3F). The highly similar phenotypes of Jasplakinolide-treated Clone 4 CTLs and TILs strongly suggest that tumor-induced disruption of actin remodeling is critical for TIL immunosuppression.

### Interface recruitment of cofilin is more sustained in TILs

T cell actin remodeling is a dynamic process, chiefly regulated by the Arp2/3 complex, which drives actin polymerization, and cofilin, which mediates the severing and depolymerisation of F-actin. To investigate mechanisms of impaired F-actin ring maintenance amongst TILs, Clone 4 CTLs and TILs expressing cofilin-GFP or Arp3-GFP were imaged (Fig. 4). Over 78% of Clone 4 CTL: Renca^+HA^ and Clone 4 TIL: Renca^+HA^ cell couples showed accumulation of cofilin-GFP in the peripheral ring distribution within 20s of tight cell coupling. In CTLs cofilin interface recruitment was highly transient. At 3 minutes of tight cell coupling only 6±3% of cell couples still displayed peripheral cofilin accumulation (Fig. 4A and E, fig. S4A). In contrast, interface accumulation of cofilin amongst TILs was sustained. At 3 minutes of tight cell coupling 55±7% of Clone 4 TIL: Renca^+HA^ cell couples still displayed peripheral cofilin accumulation (p<0.0001)(Fig. 4B and E, fig. S4B). The overall interface amount of cofilin at the immune synapse of Clone 4 TILs was also significantly (p<0.0001) higher from time point 40s onwards (Fig. 4F). To establish whether the increase in cofilin localization was associated with enhanced cofilin activation, we determined the amounts of active, i.e. non-phosphorylated, cofilin by phos-tag western blotting of Clone 4 CTL and TIL lysates activated on anti-CD3 antibody-coated plates. At all of the time points, the percentage of active, unphosphorylated cofilin was significantly (p<0.05) increased within TILs (Fig. 4H and I, fig. S4E). To establish that negative F-actin regulation in TILs is mediated by cofilin specifically, interface recruitment of another negative regulator of T cell actin dynamics, Coronin 1A, was imaged. In contrast to cofilin, interface accumulation of Coronon 1A was highly similar between CTLs and TILs (fig. S4L and M). Cofilin activation by dephosphorylation at Ser3 is controlled by multiple phosphatases. One of the key activators of cofilin in T cells is Chronophin (27). Interface recruitment of Chronophin was increased in Clone 4 TILs compared to Clone 4 CTLs, albeit at an overall low level (fig. S4F to I). Such enhanced Chronophin accumulation may contribute to increased cofilin activity. In contrast to the substantial differences in interface recruitment of cofilin between Clone 4 CTLs and TILs, the percentage of cells displaying peripheral Arp3-GFP accumulation was largely unchanged (Fig. 4C, D, and G, fig. S4C and D). As the balance between Arp2/3 complex and cofilin activity largely governs T cell actin dynamics, the prolonged cofilin interface recruitment amongst TILs is highly likely to drive the concurrent diminished maintenance of the peripheral F-actin ring.

**Fig. 4.**
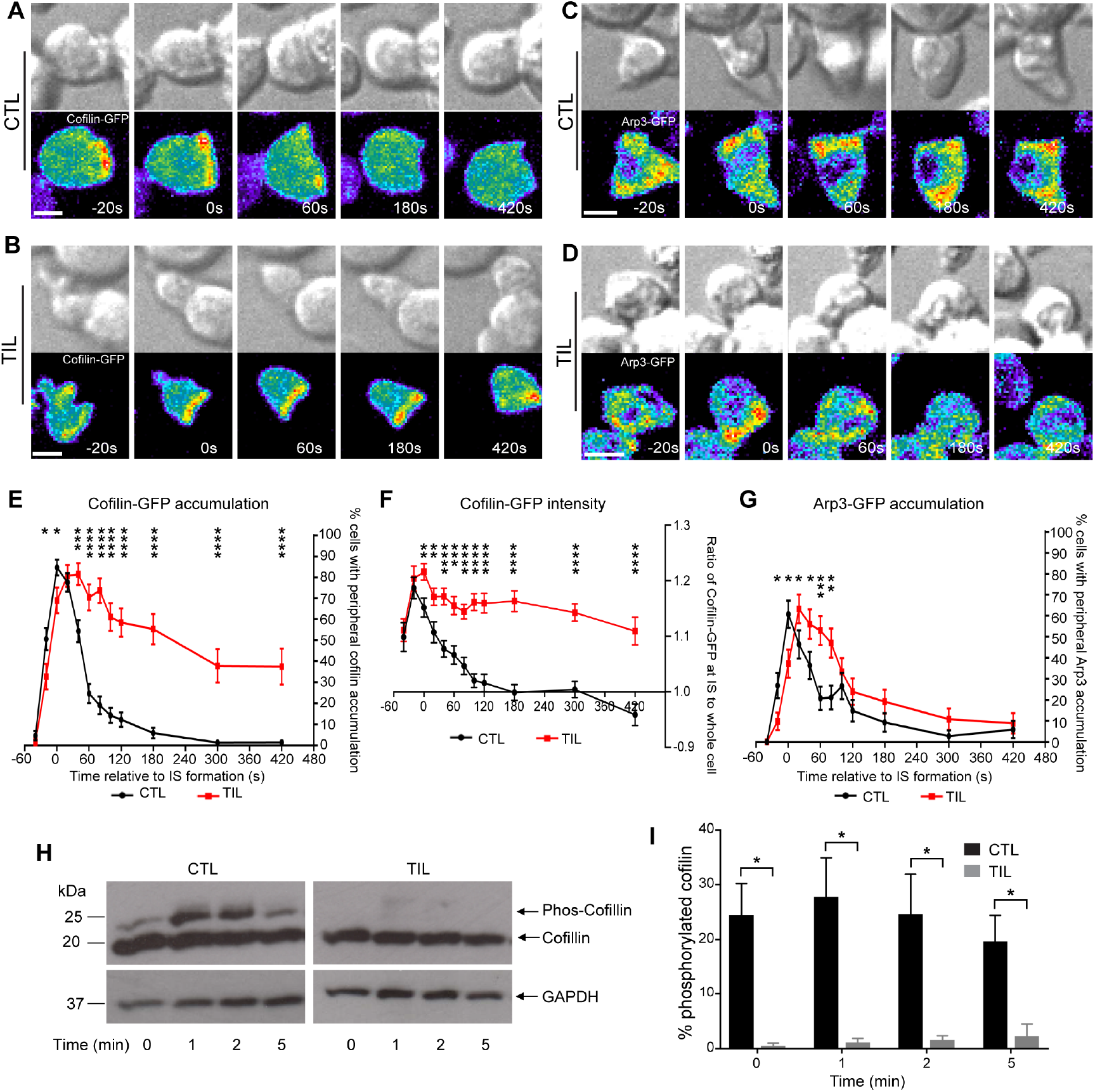
Cofilin, but not Arp2/3, displays more sustained interface accumulation in TILs. (A, B) Representative interaction over time of cofilin-GFP transduced Clone 4 CTLs (A) and TILs (B) with HA peptide pulsed Renca^+HA^ cells. Shown are matching DIC and maximum GFP projection panels with a false colour GFP intensity scale (blue to red). Scale bar shows 5μm. Complete interactions in Videos S6, S7. (C, D) Representative interaction over time of Arp3-GFP transduced Clone 4 CTLs (C) and TILs (D) with HA peptide pulsed Renca^+HA^ cells. Shown are matching DIC and maximum GFP projection panels with a false colour GFP intensity scale (blue to red). Scale bar shows 5μm. Complete interactions in Videos S10, S11. (E) Percentage over time of cofilin-GFP transduced Clone 4 CTLs and TILs displaying peripheral interface accumulation of cofilin-GFP during interaction with HA pulsed Renca^+HA^ cells. Tight cell coupling: time 0s. Mean ± SEM of 90 CTLs and 58 TILs from ≥2 separate experiments. P values calculated using proportions z test. (F) The mean fluorescence intensity of cofilin-GFP was measured across the entire T cell and at the immune synapse, defined as quarter of the T cell length adjacent to the immune synapse, for Clone 4 CTLs and TILs interacting with HA peptide pulsed Renca^+HA^ cells. The ratio of cofilin-GFP fluorescence intensity between the immune synapse and entire cell was calculated. Mean ± SEM, 47 CTLs and 56 TILs analysed from ≥2 separate experiments. P values calculated using Student’s t test. (G) Percentage over time of Arp3-GFP transduced Clone 4 CTLs and TILs displaying peripheral interface accumulation of Arp3-GFP during interaction with HA pulsed Renca^+HA^ cells. Tight cell coupling: time 0s. Mean ± SEM of 56 CTLs and 51 TILs, from ≥2 separate experiments. P values calculated using proportions z test. (H, I) Clone 4 CTLs and TILs were activated using plate bound anti-CD3 monoclonal antibody for between 1-5 minutes, as indicated, cells were lysed and the degree of cofilin phosphorylation assessed using phostag western blotting. Representative western blot (H), full blot shown in supplementary Fig. 4E. Percentage of phosphorylated cofilin over time was calculated by measuring intensities of phosphorylated and unphosphorylated cofilin bands (I). Mean ± SEM was calculated from three separate experiments. P values were calculated using Student’s t test. * p < 0.05, ** p < 0.01, *** p < 0.001, **** p < 0.0001.

In CD4^+^ T cells differences in the spatiotemporal organization of proximal T cell signaling are tightly linked to differences in actin dynamics (28-30). To similarly gain insight into signaling defects in TILs, which may underpin impaired cell couple maintenance, we investigated key elements of the spatiotemporal organization of Clone 4 CD8^+^ T cell signaling. During the activation of Clone 4 CTLs by Renca^+HA^ cells as APCs substantial μm scale clustering of TCRζ, LAT or PIP_2_ was not detected (fig. S5A to C). This is consistent with inefficient clustering of LAT and active Rac in P14 TCR transgenic CD8^+^ T cells (31) and suggests that efficient cytolytic killing does not require μm scale clustering of proximal signaling in CTLs. However, the phosphatase SHP-1 transiently accumulated at the Clone 4 Renca^+HA^ interface, almost exclusively at the interface periphery, with 60±7% of cell couples displaying SHP-1 accumulation at the time of tight cell coupling (fig. S5D to F). The frequency of interface SHP-1 accumulation in Clone 4 TILs was significantly (p≤0.05) enhanced compared to Clone 4 CTLs at multiple time points after the first minute of cell coupling. More sustained SHP-1 accumulation is consistent with a suppression of proximal T cell signaling that could diminish interface F-actin accumulation.

### Blocking PD-1 in vivo enhances TIL killing and tumor rejection

To identify potential regulators of the diminished maintenance of TIL polarization, we investigated the role of PD-1. CD8^+^ T cell PD-1 expression was upregulated in RencaHA tumors. PD-1 was expressed on 78±4% of Clone 4 TILs and 80±3% endogenous CD8^+^ TILs, significantly (p<0.01) higher than the percentage of Clone 4 CTLs expressing PD-1 (51±9%)(Fig. 5A). The mean fluorescence intensity of the PD-1 staining was also elevated on TILs (fig. S6A). To investigate whether increased PD-1 expression contributed to Clone 4 TIL dysfunction, RencaHA tumor-bearing mice were treated with anti-PD-1 blocking monoclonal antibody (mAb) in combination with adoptive transfer of Clone 4 CTLs (Fig. 5B). Over the six-day treatment period, tumor growth was significantly (p=<0.01) lower in the anti-PD-1 antibody-treated mice (62±74% increase in tumor volume) as compared to buffer only or isotype-treated control mice (166±123% increase in tumor volume)(Fig. 5C). In addition, killing of Renca^+HA^ tumor cells ex vivo by Clone 4 TILs isolated from anti-PD-1 treated mice was significantly (p<0.05) enhanced (Fig. 5D) but did not reach levels achieved by CTLs (Fig. 3F).

**Fig. 5.**
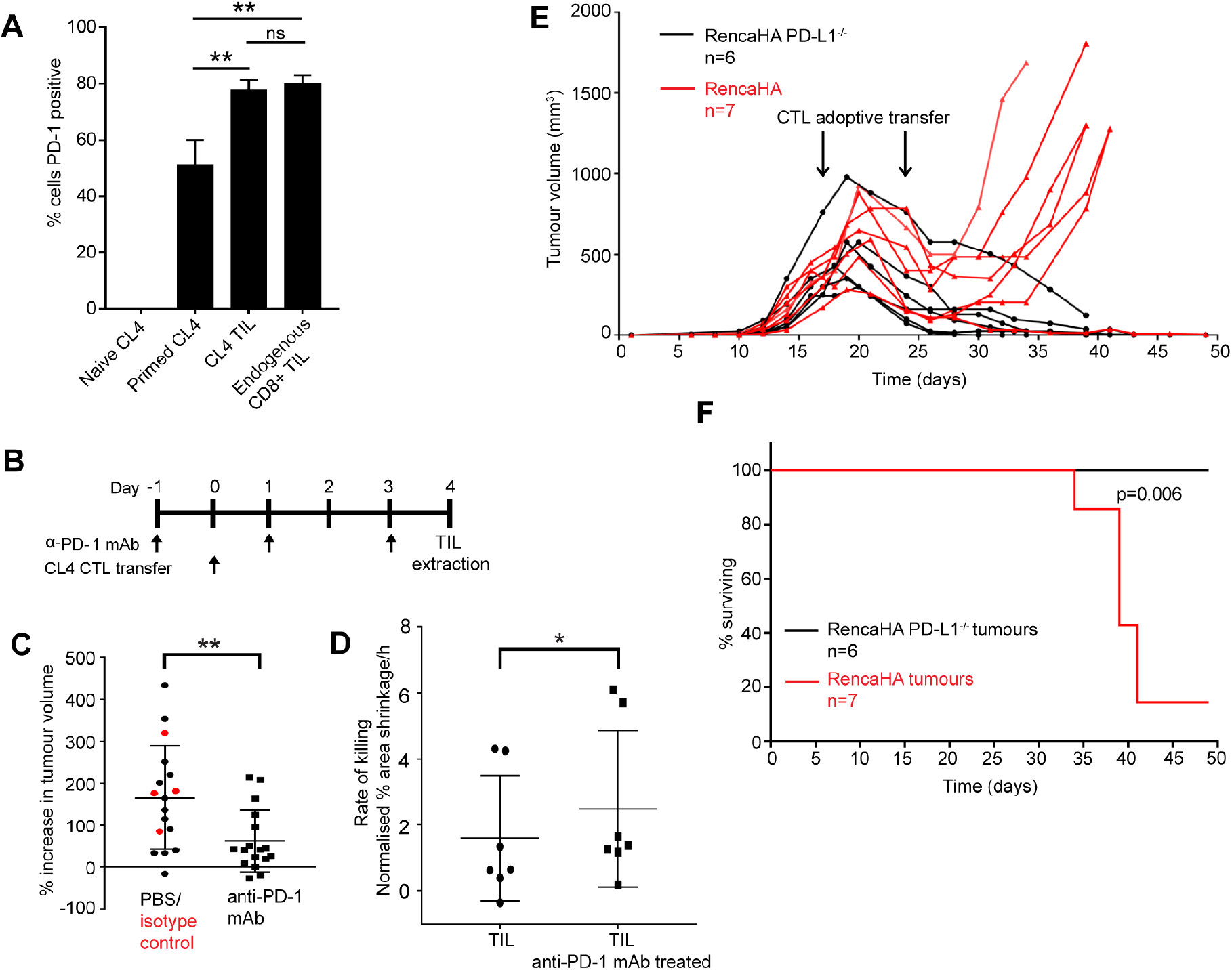
Loss of PD-1 signalling in vivo, but not ex vivo, improves TIL killing ability. (A) Naïve and primed Clone 4 CTLs, Clone 4 TILs and endogenous CD8^+^ TILs were stained for PD-1 expression and analysed by flow cytometry, using PD-1 FMO cells for gating. The percentage of cells expressing PD-1 is shown. Data from 4 separate experiments, involving 16 tumours. Mean ± SEM. (B) Timeline of anti-PD-1 mAb and Clone 4 CTL adoptive transfer immunotherapy strategy. Injection of anti-PD-1 and CTLs both done via i.v. injection. (C) Mice were inoculated s.c. with 1×10^6^ RencaHA cells. Mice with approximately equal sized tumours (approx. 500mm^3^) were given anti-PD-1 mAb (or control) and Clone 4 CTLs as detailed in (B). Tumour volume was recorded at the beginning and end of treatment. Percentage change in tumour volume was calculated. Control mice received either PBS or isotype control mAb (red points). Data shows mean ± SD, 17 mice per condition over 5 separate experiments. (D) Average rate of CTV labelled, HA peptide pulsed Renca^+HA^ cell death following incubation with Clone 4 TILs from RencaHA tumour bearing mice receiving anti-PD-1 mAb (or PBS control) and Clone 4 CTL adoptive transfer as detailed in (B). Rate calculated as percentage Renca^+HA^ density decrease per hour. Tumour volumes were measured prior to treatment inititation and ranged from 300-600mm^3^. Mice were paired based on matched tumour volume at the start of treatment. Data shows mean ± SD. (E) Volume of individual tumours formed by RencaHA or RencaHA PD-L1^-/-^ cells following s.c. inoculation. Mice received 2×10^6^ Clone 4 CTL by i.v. injection on days 17 and 24. Data pooled from two separate experiments, n=7 RencaHA and n=6 RencaHA PD-L1^-/-^ tumour bearing mice. (F) Kaplan-Meier survival curve for RencaHA and RencaHA PD-L1^-/-^ tumour bearing mice from (E). * p < 0.05, ** p < 0.01, *** p < 0.001, **** p < 0.0001; p values calculated using one-way ANOVA (A, G), Student’s t test (C), paired Student’s t test (D), and Mantel-Cox test (F).

PD-1 is engaged by its ligand PD-L1 as expressed on tumor cells and/or tumor-associated immune cells. To investigate the functional relevance of tumor-expressed PD-L1, a RencaHA-PD-L1^-/-^ cell line was generated (fig. S6B). Loss of PD-L1 expression induced spontaneous tumor regression in 4/10 mice, and significantly (p=<0.01) increased survival (fig. S6C and D). Upon two adoptive transfers of Clone 4 CTLs, 6/6 mice bearing RencaHA-PD-L1^-/-^ tumors underwent complete tumor clearance, compared to only 1/7 control mice (Fig. 5E and F)(p<0.01), establishing that PD-L1 on RencaHA cells greatly contributes to suppression of adoptively transferred Clone 4 CTLs. In combination experiments with blocking anti-PD-1 antibodies and PD-L1-deficient tumor cells establish that PD-1 is a critical element of in vivo tumor immune suppression in the RenaHA model.

### PD-1 promotes the impaired maintenance of TIL polarization in vivo

To investigate whether blockade of PD-1 engagement within the tumor microenvironment restores TIL polarization, we studied TILs isolated from RenacHA tumor-bearing mice treated with anti-PD-1 mAb or from RencaHA-PD-L1^-/-^ -bearing mice ex vivo. The percentage of TILs with translocation of at least one interface diameter over the target cell surface following immune synapse formation was significantly (p=<0.0001) reduced (Fig. 6A) upon treatment with anti-PD-1 mAb or use of RencaHA-PD-L1^-/-^ tumors to 9±0.7% and 11±0.4%, respectively, as compared to 14±0.2% in buffer only-treated TILs. This reduction suggests a partial restoration of cell couple stability. The production of off-synapse lamellae was not reduced by PD-1 blockade (fig. S7A and B).

**Fig. 6.**
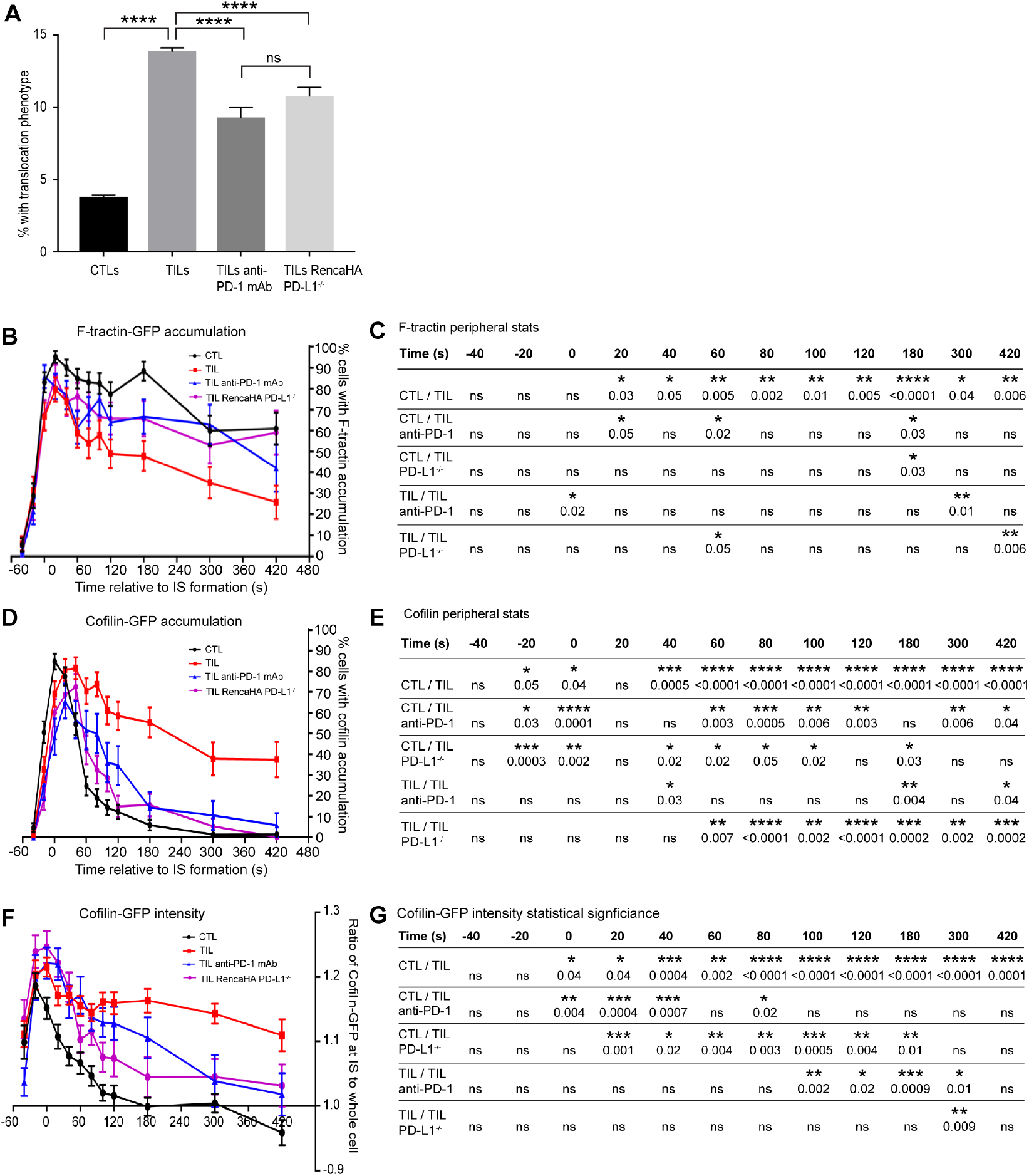
Loss of PD-1 signalling in vivo significantly rescues normal F-actin and cofilin regulation. (A) Clone 4 TILs were isolated from RencaHA tumour bearing mice treated with anti-PD-1 mAb, as in Fig. 5B, and from RencaHA-PD-L1^-/-^ tumour bearing mice. The percentage of cells with translocation of more than one interface diameter around the HA peptide pulsed RencaHA^+HA^ target cell is given in comparison to Clone 4 CTLs and TILs from RencaHA tumour bearing mice (data shown previously, Fig. 2H). Mean ± SEM, 132 CTL, 137 TIL, 54 TIL from anti-PD-1 treated mice, 65 TIL from RencaHA-PD-L1^-/-^ tumour bearing mice. Data from ≥3 separate experiments per condition. (B, C) The percentage over time of F-tractin-GFP transduced Clone 4 cells displaying peripheral F-tractin-GFP interface accumulation during interaction with HA peptide pulsed Renca^+HA^ target cells. Data is shown for Clone 4 CTLs and TILs (data shown previously, Fig. 3D) as well as TILs from RencaHA tumour bearing mice treated with anti-PD-1 mAb, as in Fig. 5B, and TILs from RencaHA-PD-L1^-/-^ bearing mice. Tight cell coupling: time 0s. Mean ± SEM, 60 CTLs, 51 TILs, 42 TILs from anti-PD-1 treated mice, 43 TIL from RencaHA-PD-L1^-/-^ bearing mice. Data from ≥2 separate experiments per condition. P values calculated by proportions z test (C). (D, E) The percentage over time of cofilin-GFP transduced Clone 4 cells displaying peripheral cofilin-GFP interface accumulation during interaction with HA peptide pulsed Renca^+HA^ target cells. Data is shown for Clone 4 CTLs and TILs (data shown previously, Fig. 4E) as well as TILs from RencaHA tumour bearing mice treated with anti-PD-1 mAb, as in Fig. 5B, and TILs from RencaHA-PD-L1^-/-^ bearing mice. Tight cell coupling: time 0s. Mean ≥ SEM, 90 CTLs, 58 TILs, 32 TILs from anti-PD-1 treated mice, 55 TIL from RencaHA-PD-L1^-/-^ bearing mice. Data from ≥2 separate experiments per condition. P values calculated by proportions z test (E). (F, G) The mean fluorescence intensity of cofilin-GFP was measured across the entire T cell and at the immune synapse, defined as quarter of the T cell length adjacent to the immune synapse, for Clone 4 CTLs, TILs, TILs from RencaHA tumour bearing mice treated with anti-PD-1 mAb, as in Fig. 5B, and TILs from RencaHA-PD-L1^-/-^ bearing mice, during interaction with HA peptide pulsed Renca^+HA^ cells. The ratio of cofilin-GFP fluorescence intensity between the immune synapse and entire cell was calculated. Tight cell coupling: time 0s. Mean ± SEM, 47 CTLs, 56 TILs, 29 TILs from anti-PD-1 treated mice, 36 TILs from RencaHA-PD-L1^-/-^ bearing mice. Data from ≥2 separate experiments. p values calculated using Student’s t test (G). (H,I) The percentage over time of F-tractin-GFP transduced Clone 4 CTLs and TILs, with and without acute in vitro anti-PD-1 mAb treatment, displaying peripheral F-tractin-GFP interface accumulation during interaction with HA peptide pulsed Renca^+HA^ cells. Tight cell coupling: time 0s. Mean ± SEM, 60 CTLs (data shown previously in Fig. 3D), 41 CTL treated with anti-PD-1 mAb, 51 TILs (data shown previously in Fig. 3D), 28 TILs treated with anti-PD-1 mAb. All data from ≥2 separate experiments. Corresponding P values were calculated using proportions z test (I). * p < 0.05, ** p < 0.01, *** p < 0.001, **** p < 0.0001; p values calculated using one-way ANOVA (A, J), proportions z-test (C and E) or Student’s t test (G).

Next we investigated F-actin as a key regulator of TIL morphology. In TILs after in vivo anti-PD-1 mAb treatment or from RencaHA-PD-L1^-/-^ tumor-bearing mice the percentage of TILs displaying peripheral F-tractin-GFP accumulation (Fig. 6B and C, fig. S7C and D) was increased 3 minutes after tight cell couple formation and later. For example, while at 5 minutes after tight cell coupling only 35±8% of Clone 4 TILs displayed peripheral F-tractin accumulation 63±9% of Clone 4 TILs from anti-PD-1-treated mice did (p=0.01)(Fig. 6C). The amount of F-tractin recruitment to the immune synapse was not significantly altered (fig. S7G). Next we investigated cofilin as a critical F-actin regulator in TILs. The percentage of TILs displaying peripheral cofilin-GFP was greatly reduced at later time points upon in vivo treatment with anti-PD-1 mAb or in TILs from RencaHA-PD-L1^-/-^ tumor-bearing mice (Fig. 6D and E). For example, while the percentage of Clone 4 TILs with peripheral cofilin-GFP accumulation did not drop below 38±9% at any time point, it did not exceed 15±5% at any point after 2min of cell coupling when using Clone 4 TILs from RencaHA-PD-L1^-/-^ tumor-bearing mice (p≤0.002 at every time point). The amount of cofilin-GFP at the immune synapse was also reduced in the TILs from tumors with blocked PD-1 engagement (Fig. 6F and G). For example at 5min after tight cell coupling, cofilin-GFP was enriched 1.1±0.02-fold at the cellular interface in control Clone 4 TILs while such enrichment was significantly (p≤0.01) reduced to 1.0±0.04-fold and 1.0±0.03-fold upon in vivo treatment with anti-PD-1 mAb or in TILs from RencaHA-PD-L1^-/-^ tumor-bearing mice, respectively. In combination the changes in morphology, F-actin and cofilin establish that sustained PD-1 engagement in the tumor microenvironment is critical for the induction of the polarization-impaired state of Clone 4 TILs.

### Acute in vitro blockade of PD-1 does not restore killing and cytoskeletal polarization

To further understand the role of PD-1 in regulating target cell killing and cytoskeletal polarization, we investigated whether PD-1 blockade just during the in vitro killing or imaging assays (‘acute’) regulates target cell lysis. When Clone 4 TILs or CTLs were treated with the same anti-PD-1 antibody as used in vivo or when RencaHA-PD-L1^-/-^ target cells were used target cell killing was not significantly altered (Fig. 7A). In addition, Clone 4 TILs or CTLs also did not show altered killing of RencaHA target cells overexpressing PD-L1 (Fig. 7A, fig. S6E). PD-1 thus did not acutely regulate cytolysis by TILs and CTLs in vitro.

**Fig. 7.**
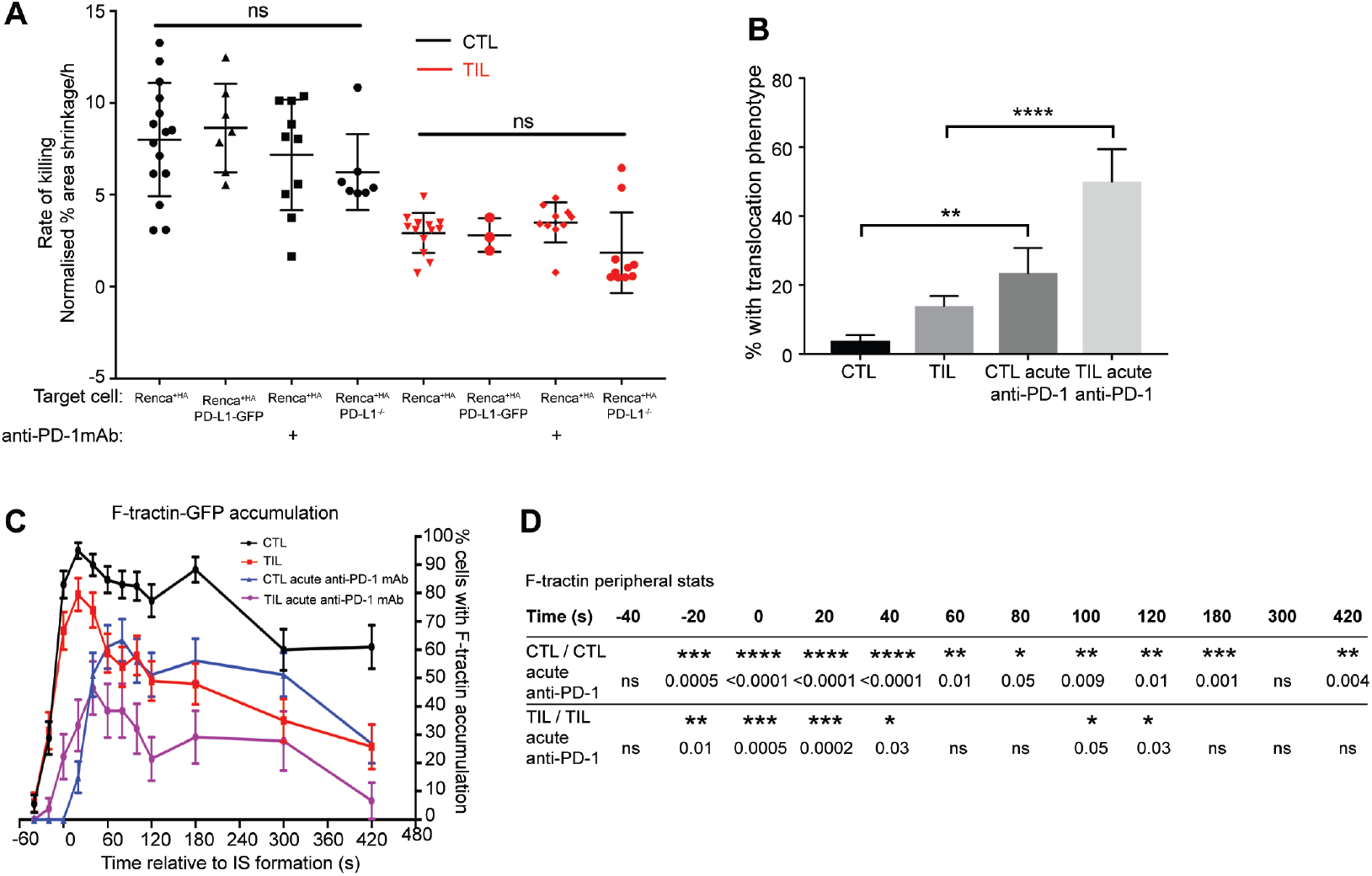
Acute in vitro blockade of PD-1 does not restore killing and cytoskeletal polarization. (A) Average rate of CTV labelled, HA peptide pulsed Renca^+HA^, Renca^+HA^-PD-L1-GFP over-expressing, or Renca^+HA^-PD-L1^-/-^ cell death by Clone 4 CTLs (black) or TILs (red), with or without in vitro anti-PD-1 treatment. Rate of death calculated as percentage Renca density decrease per hour. T cells receiving anti-PD-1 treatment in vitro were incubated with anti-PD-1 mAb for 1 hour prior to and during the killing assay. Data shows mean ± SD from a minimum of three separate experiments. Data for CTL with Renca^+HA^ and TIL with Renca^+HA^ previously shown in Fig. 1D. (B) Clone 4 CTLs and TILs, with and without acute in vitro anti-PD-1 treatment, coupled to HA peptide pulsed Renca^+HA^ cells, were analysed over time to determine the percentage of T cells displaying translocation as defined as the migration of the T cell further than one immune synapse diameter. Data shows mean ± SEM. Translocation data for CTLs and TILs without anti-PD-1 treatment was previously shown in Fig. 2H. (C,D) The percentage over time of F-tractin-GFP transduced Clone 4 CTLs and TILs, with and without acute in vitro anti-PD-1 mAb treatment, displaying peripheral F-tractin-GFP interface accumulation during interaction with HA peptide pulsed Renca^+HA^ cells. Tight cell coupling: time 0s. Mean ± SEM, 60 CTLs (data shown previously in Fig. 3D), 41 CTL treated with anti-PD-1 mAb, 51 TILs (data shown previously in Fig. 3D), 28 TILs treated with anti-PD-1 mAb. All data from ≥2 separate experiments. * p < 0.05, ** p < 0.01, *** p < 0.001, **** p < 0.0001; p values calculated using one-way ANOVA (A, B) or proportions z-test (D)

Next we investigated the effect of acute PD-1 blockade on CD8^+^ T cell morphology and F-actin distributions as critical elements of cellular polarization (Fig. 7B to D, fig. S8). Upon acute treatment with anti-PD-1 mAb the percentage of Clone 4 CTL and TIL cell couples with Renca^+HA^ target cells that showed translocation of the T cell from its initial site of target cell binding by at last one interface diameter increased from 4±2% of CTLs to 24±7% (p<0.01) and from 14±3% of TILs to 50±9% (P<0.0001)(Fig. 7B), respectively. In addition, acute PD-1 blockade substantially delayed the formation of a peripheral actin ring. Whilst at the time of tight cell coupling with Renca^+HA^ target cells (t=0s) 83±5% of Clone 4 CTLs had already formed a peripheral F-actin ring, no Clone 4 CTL did so upon anti-PD-1 mAb treatment (p<0.0001)(Fig. 7C and D). Similarly, whilst 62±7% of Clone 4 TILs had already formed a peripheral F-actin ring, only 22±8% did so upon anti-PD-1 mAb treatment (p=0.0005). While PD-1 blockade in vivo thus restored a substantial fraction of the impaired cell couple maintenance of TILs acute in vitro blockade could not do so. On the contrary, some elements of cell polarity were actually impaired.

## Discussion

The mechanisms underpinning the defective killing ability of TILs are still uncertain. A number of alterations in TIL activation have been described, including less actin polymerization (32-34), increased expressing of inhibitory signaling mediators such as SHP-1 and Cish (35, 36) and decreased functional avidity of the T cell receptor (37). Metabolic defects in the glucose-poor tumor microenvironment are likely as enhancement of TIL glycolysis leads to improved tumor killing (38, 39). By establishing an experimental system for the rapid ex vivo imaging of the antigen-driven interaction of TILs with their tumor target cells we have discovered a cellular mechanism of tumor immune suppression: Impaired maintenance of cellular polarization diminished TIL killing ability as regulated in vivo by the inhibitory receptor PD-1 (Fig. 8). Time is a critical element of this mechanism as, first, the polarization defect only began to appear two minutes after tight cell coupling and, second, PD-1 was only effective over days within the tumor microenvironment but not acutely.

**Fig. 8.**
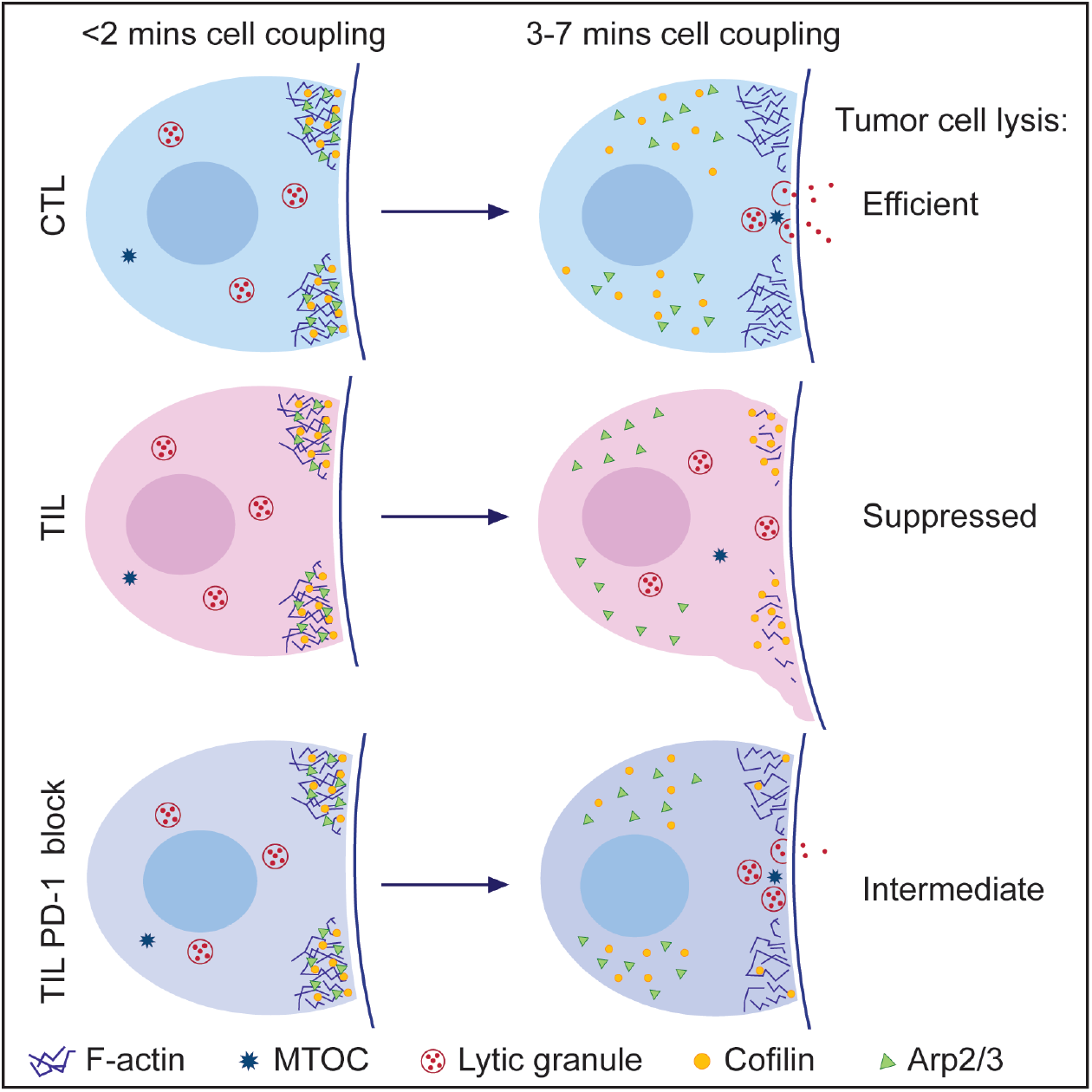
Graphical summary.

TILs could efficiently form cell couples with their tumor target cells, yet 1-2 minutes into cell coupling TILs increasingly lost the peripheral F-actin ring and developed lamellae pointing away from the target cell interface. While recruitment of the Arp2/3 complex to the interface periphery as the principal means of T cells to generate F-actin (40) was similar between TILs and CTLs, recruitment of cofilin as a principal F-actin severing protein differed substantially. It was highly transient in CTLs yet much more sustained in TILs. This was a specific defect as recruitment of Coronin1A as another negative regulator of actin dynamics was much more similar. Our data are consistent with a time-dependent switch between two principal roles of cofilin, promotion of dynamic F-actin turnover and F-actin disassembly. Cofilin can drive efficient F-actin turnover by providing actin monomers and mother filaments for actin polymerization for example in cell motility and cytokinesis (41-43). The coincident recruitment of cofilin and Arp3 to the T cell tumor target cell interface and generation of a peripheral F-actin ring during the first minute of cell coupling is consistent with the F-actin promoting role of cofilin. This constructive role of cofilin is consistent with the established need of cofilin for the enhancement of T cell activation by costimulation (44). Upon reduced F-actin generation cofilin disassembles F-actin structures. Accelerated loss of the peripheral F-actin ring in TILs starting 1-2 minutes after tight cell coupling is therefore consistent with the more sustained interface accumulation of cofilin. Four elements of tumor immune suppression have already been linked to cofilin supporting the proposed role of enhanced cofilin activity in diminished TIL function. The prostaglandin PGE_2_ is an established element of Clone 4 tumor immunosuppression (45) and inhibits actin polymerization in phagocytosis through activation of cofilin (46). Similarly, adenosine receptors that in the tumor microenvironment mediate immune suppression by adenosine (6, 7) inhibit cofilin inactivation in cardiomyocytes (47). Cofilin activation can also be inhibited by oxidation in T cells and mesenchymal cell motility (48, 49). The low oxygen environment of solid tumors (50) thus is consistent with enhanced cofilin activity. Finally, low levels of intracellular ATP induce cofilin activation by Chronophin in neurons (51), a process that is equally conceivable in the energy-starved tumor microenvironment.

We observed that diminished PD-1 engagement in the tumor could partially restore the ability of TILs to maintain a polarized cell couple with the tumor target cells and kill. In intriguing contrast, acute blockade of PD-1 during its in vitro interaction with Renca tumor target cells did not rescue impaired TIL polarization nor killing. On the contrary, Clone 4 CTL polarization was impaired upon acute blockade of PD-1 as to be further investigated in the future. These data suggest that extended in vivo engagement of PD-1 induces a ‘polarization-impaired state’ that requires hours to days to form and revert. In support, 24h co-culture of primary T cells with various tumor cell types diminishes the ability of the T cells to form cell couples with allogeneic antigen-pulsed B cells thereafter (33) as partially controlled by PD-1 (32). Providing estimates of the time scale involved in switching T cells between suppressed and active states, in vitro culture of TILs in IL-2 for more than 6h can overcome diminished TIL killing of tumor target cells (52) and in vitro resting of TILs for >24h restores impaired tetramer binding to the TCR (37).

At this time it remains unresolved what is more important in the establishment of the TIL polarization-impaired state, the extended time of PD-1 engagement in the tumor microenvironment or synergy of PD-1 with other suppressive factors provided by the tumor. In addition, molecular changes underpinning the TIL polarization-impaired state remain to be discovered. The need for extended PD-1 engagement within the context of the tumor microenvironment makes it difficult to link established signaling roles of PD-1, that are largely investigated acutely and in vitro, to the in vivo regulation of TIL polarization and function by PD-1. Nevertheless, transcriptional changes in TILs are conceivable, as sustained engagement of another receptor, the TCR by antigen, can substantially alter the transcriptional program of T cells (53) as also shown in the context of cancer (54). Metabolic changes are conceivable, as enhanced tumor killing by blockade of PD-1 is associated with increased glycolysis (39). In addition, engagement of the costimulatory receptor CD28 is required for the restoration of T cell function by PD-1 blockade (55, 56). While Clone 4 TILs may have access to the CD28 ligands CD80 and CD86 on the surface of tumor infiltrating professional antigen presenting cells within the tumor microenvironment, there is no such access during the in vitro interaction of Clone 4 T cells with Renca^+HA^ target cells. Future investigations of the mechanisms of PD-1 function will have to take into account the difference between sustained in vivo and acute in vitro roles established here.

## Materials and Methods

### Cells and Media

‘Complete medium’ consisted of RPMI-1640 plus L-glutamine (Gibco) supplemented with 10% FBS (Hyclone defined, US source, GE healthcare), 50μM β-Mercaptoethanol (Gibco), and PenStrep (Gibco) at 100U/mL penicillin and 100μg/mL streptomycin. RencaWT cells were maintained in complete medium, whilst RencaHA cells were maintained in complete medium supplemented with 100ug/ml geneticin. Clone 4 T cells were maintained in complete medium supplemented with 50U/mL rh-IL-2 (NIH/NCI BRB preclinical repository)(‘IL-2 medium’). Phoenix cells were maintained in ‘Phoenix incomplete medium’ consisting of DMEM with 4.5g/L D-glucose, L-glutamine and sodium pyruvate (Gibco) supplemented with 10% FBS (Hyclone define, US source, GE healthcare), MEM non-essential amino acids (Gibco), and PenStrep (Gibco) at 100U/mL penicillin and 100μg/mL streptomycin. Long terms Phoenix cell stocks were kept in ‘Phoenix complete medium’, which is Phoenix incomplete medium supplemented with 300μg/mL Hygromycin (Invitrogen) and 1μg/mL Diptheria toxin (Sigma).

### Mice

Clone 4 TCR transgenic mice (RRID: IMSR_JAX:005307) were bred and maintained at the University of Bristol under specific pathogen free conditions, with unlimited access to water and standard chow. Clone 4 offspring were phenotyped by staining PBMCs with anti-CD8-APC (53-6.7, Biolegend, RRID: AB_312750) and anti-Vβ8.2 FITC (KJ16-133, eBioscience, RRID: AB_465261) antibodies. All Clone 4 mice were culled for experimental use between 6 and 12 weeks of age. BALB/c mice were purchased from Charles River Laboratories at six weeks of age and then maintained at the University of Bristol under specific pathogen free conditions. All mice were culled using schedule one methods and any experimental procedures were conducted in accordance with protocols approved by the UK Home Office.

### Clone 4 CD8+ T cell isolation and stimulation

Procedures used for the isolation and stimulation of TCR transgenic T cells have been extensively detailed elsewhere (22, 29, 57). Briefly, red blood cells were removed from dissociated spleens using ACK lysis buffer (Gibco) and splenocytes were plated at a density of 5×10^6^ cells per well, in 1 mL of complete medium, in a 24-well plate. K^d^HA peptide (IYSTVASSL) was added to a final concentration of 1μg/mL and cells were incubated overnight. Primed cultures were stringently washed five times in PBS to remove unbound peptide and then retrovirally transduced (detailed below). Following transduction, cells were re-plated at 4×10^6^ cells per well, in 2 mL IL-2 medium, in a 24-well plate. Cells were cultured for a further two days. CTLs were sorted and imaged, or adoptively transferred into RencaHA tumor bearing mice, on day four.

### T cell retroviral transduction

Procedures and plasmids used for retroviral transduction of TCR transgenic T cells have been extensively detailed elsewhere (22, 29, 57). Briefly, transfection of a GFP-sensor expressing plasmid was achieved using calcium phosphate precipitation. After transfection cells were cultured for a further 48 hours, then the retrovirus-containing medium was collected and used to transduce T cells. Clone 4 splenocytes were activated with K^d^HA peptide for 24 hours, as above. After washing, each well of cells was resuspended in 2 mL retrovirus-enriched supernatant from transfected Phoenix-E cells. Protamine sulphate was added to a final concentration of 8μg/mL. Plates were centrifuged for 2 hours at 200xg, at 32°C. Following centrifugation, supernatant was carefully removed without disturbing the T cell layer and fresh IL-2 medium was added. Cells were cultured as above.

### RencaHA tumor growth in BALB/c mice

To induce tumor growth, female 6-8-week old BALB/c mice were injected subcutaneously with 1×10^6^ RencaHA cells suspended in PBS. Injections were performed in the scruff of the neck. Mice were monitored regularly following injection to assess tumor growth. Tumors were measured using calipers and calculated using the equation: volume (V)=L2xW/2, where length (L) is the longest measurement and width (W) is measured perpendicularly to length. Mice were culled if tumor volume exceeded 1500mm^3^ or any humane end points were reached.

### Adoptive transfer of Clone 4 CD8+ T cells

The adoptive transfer of Clone 4 CTLs was performed via tail vein injection. CTLs were collected from culture, and sorted to isolate GFP-positive cells, as detailed below. Cells were resuspended in PBS at a concentration of 2×10^7^ cells per mL, and 200μL injected into the tail vein of a tumor bearing mouse, giving a total of 4×10^6^ T cells per treatment. Unless otherwise specified, only mice bearing tumors over 500μm^3^ were used and tumors were harvested 96 hours after transfer.

### In vivo anti-PD-1 immunotherapy

Mice received three doses of either anti-PD-1 (RMP1-14, Bio X cell, RRID: AB_10949053) or rat IgG2a isotype control (2A3, Bio X cell, RRID: AB_1107769) monoclonal antibodies on days one, three and five of treatment. CTL adoptive transfer occurred on day two and mice were culled on day six. Tumor measurements were taken on day one and six, prior to culling. Each dose consisted of 250μg mAb in 200μL PBS, and was given via tail vein injection. In the event that a tail vein injection could not be performed, antibody was given intraperitoneally.

### Isolation of TILs

Tumors were collected into 3.2mL RPMI-1640 (no FBS), roughly chopped with sterile scissors, and tumor cell dissociation enzymes (Miltenyi Biotec) were added as per manufacturers instructions. Tumors were incubated at 37°C for 45 minutes, and briefly vortexed every 10 minutes to enhance dissociation. The tumor suspension was then passed through a 40μm sieve with RPMI-1640, and red blood cells removed using ACK lysis buffer. Cells were pelleted, then resuspended in 600μL MACs buffer (PBS (no Ca^2+^ or Mg^2+^), 2mM EDTA, 0.5% BSA) with 65μL MACs microbeads (Miltenyi) and incubated at room temperature for 30 minutes. CD8a (Ly-2) microbeads were used to enrich for CD8^+^ T cells, whilst CD45 microbeads were used to enrich for all lymphocytes. Positive selection of the magnetically labeled cell populations was performed using MACS LS-columns (Miltenyi Biotec). Collected cells were pelleted and resuspended in 500μL imaging buffer, for immediate sorting based on GFP signal, or 500μL FACS buffer for antibody staining.

### Imaging of Clone 4 CTLs and TILs

Approximately 72 hours after retroviral transduction, or immediately after TIL extraction, T cell cultures were resuspended in ‘imaging buffer’ (10% FBS in PBS with 1mM CaCl_2_ and 0.5mM MgCl_2_) plus propidium iodide (PI). A BD Influx cell sorter (BD Bioscience) was used to isolate GFP-positive T cells (22). As target cells, 1×10^6^ RencaWT cells were pulsed with K^d^HA peptide at a final concentration of 2μg/mL for 1 hour. These pulsed RencaWT cells will be referred to as Renca^+HA^ cells. Glass bottomed, 384-well optical imaging plates (Brooks life science systems) were used for all imaging experiments. Imaging was done at 37°C using a Perkin Elmer UltraVIEW ERS 6FE confocal system attached to a Leica DM I6000 inverted epifluorescence microscope and a Yokogawa CSU22 spinning disk. A 40x oil-immersion lens (NA=1.25) was used for all imaging experiments, unless otherwise stated. Prior to imaging, 50μL of imaging buffer was added to the well, followed by 5μL of T cells. T cells were allowed to settle for several minutes. Once a monolayer of T cells had formed, 4μL of Renca^+HA^ cells were carefully added to the top of the well. Images were acquired for 15 minutes. Every 20 seconds, a z-stack of 21 GFP images (1μm z-spacing) was acquired, as well as a single, mid-plane differential interference contrast (DIC) reference image. After imaging, GFP-files were saved as individual z-stacks for each time point, and both DIC and GFP files were exported as TIFF files for subsequent analysis.

Some imaging experiments involved prior treatment of T cells with 40nM Jasplakinolideor or 10μg/mL anti-PD-1 mAb (RMP1-14, Bio X cell, RRID: AB_10949053). In both cases, reagent was added at the desired concentration for 30 minutes prior to imaging, and cells were incubated at 37°^C^. In both cases, the reagent was also present in the well throughout imaging.

### Analysis of live cell imaging data to assess spatiotemporal patterning

Image analysis protocols have previously been described (22, 29, 57). DIC files and GFP z-stacks were imported into Metamorph image analysis software (Molecular Devices). For each imaging time point, the GFP z-stack was assembled into a maximum projection. CTLs forming conjugates with RencaWT^HA^ target cells were identified and the time point at which a tight-cell couple forms was assessed using the DIC reference images. Tight cell couple formation was defined as the first time point at which a maximally spread immune synapse forms, or two frames following initial cell contact, whichever occurred first. Time points -40 to 120, 180, 300 and 420 seconds were analyzed, giving a total of 12 data points for each cell couple, with an emphasis on events occurring shortly after immune synapse formation. For each time point, the 3D GFP data was used to assess protein accumulation patterns at the immune synapse, and the cell was sorted into one of seven patterning categories for that time point. These pattern classifications have been extensively described in previous publications (22, 30). For each condition, the percentage of cells displaying each pattern at each time point was calculated. To assess CTL and TIL morphology, every DIC frame following tight cell couple formation was assessed for the presence of off-synapse lamella, defined as a transient membrane protrusion pointing away from the immune synapse, followed by retraction. For each cell couple, the initial position of the immune synapse on the Renca^+HA^ target cell was compared to the position in the final frame. If the T cell had migrated by more than the immune synapse diameter, this was classed as significant translocation.

### T cell killing assays

Renca^+HA^ or Renca^WT^ cells were stained with 10μM cell trace violet (CTV) and plated at a density of 1×10^4^ cells per well of a 384-well plate (Perkin Elmer) in 100μL complete medium. Cells were incubated at 37°C for a minimum of five hours to allow adherence and spreading. Prior to imaging, medium was exchanged for 50μL/well imaging buffer. Clone 4 CTLs or TILs were resuspended in imaging buffer at a density of 2×10^5^ cells/mL. The imaging plate was then mounted on a Leica DMI6000 inverted epifluorescence widefield microscope, fitted with a 37°C incubation chamber, a humidified CO_2_ enrichment attachment, a motorized stage, and adaptive focus control enabled image analysis software (Leica LAS-X). Using the multi-site image acquisition software, the DAPI channel was used to image the CTV stained Renca cells and the positions for 2-3 fields of view were saved per well. Once positions had been saved, 50μL of the T cell culture (giving a total of 1×10^4^ cells/well) were added per well. After T cells were added, images of CTV-stained Renca cells were acquired every 30 minutes for 10 hours, using adaptive focus control to prevent drift. Images were analyzed as a separate.tiff movie for each saved field of view, using Metamoph image analysis software. A mask was generated using thresholding for light objects and adjusted to match the fluorescent signal. The same thresholding criteria were then used for all conditions within that experiment. To assess killing ability of T cells, the area of the field covered by CTV-stained Renca^+HA^ cells was measured. For each condition, the average percentage change in cell area was calculated for each time point, and a gradient describing the rate of cell area change was calculated. This was corrected for a control condition, in which no T cells were added and the RencaWT^HA^ cells proliferated, providing a normalized value for the rate of cell area change across all experiments.

### Generation of RencaHA-PD-L1-/-cell line

Methods used closely follow those described in (58). SgRNAs were designed using three separate programmes: Benchling, Broad institute, and MIT CRISPR design. Results were compared and six top common hits were selected. DNA sequences encoding these sgRNAs were designed as oligonucleotides which could be inserted into the pSpCas9(BB)-2A-GFP vector (Addgene). The primer sequences ultimately used to produce the selected B10 clone are detailed in the table below. Vectors were transfected into RencaHA cells using Lipofectamine 2000 as per manufacturer’s instructions. After 24 hours, FACS was used to separate single-cell GFP-positive clones into separate wells of a 96-well plate. Colonies were expanded and screened for PD-L1 loss by staining and flow cytometry. Several clones were further screened to compare MHC class I expression, HA expression and in vitro proliferative ability to the original RencaHA cell line, and the most similar clone (B10) was selected.

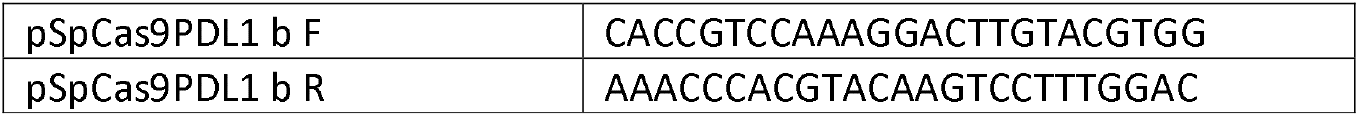

Primers encoding gRNA sequences for CRISPR/Cas9 mediated knockout of PD-L1.

### Generation of RencaWT PD-L1-GFP over expressing cells

The gene encoding PD-L1 was cloned into a pSRαGFP vector expressing GFP and geneticin resistance, to create a PD-L1-GFP fusion construct. This vector was transfected into RencaWT cells using lipofectamine 2000 (Invitrogen), as per the manufacturer’s instructions. Cells were cultured without antibiotic for 48 hours, then exchanged into 200μg/mL geneticin for 48 hours to kill untransfected cells. Cells were then washed and exchanged into complete medium with 100μg/mL geneticin, for maintenance of the new cells. Cells were checked under a widefield microscope for expression and membrane localization of PD-L1-GFP. Cells were expanded, before FACS was performed to select for GFP-positive cells. Successfully transduced cells were cultured for a further two weeks, then analyzed by anti-PD-L1 antibody staining and flow cytometry to assess for over expression of GFP-conjugated PD-L1.

### Phos-tag western blotting to assess cofilin phosphorylation

Round-bottom, 96-well plates were coated overnight with 10μg/mL anti-CD3 antibody (145-2C11, BioLegend, RRID: AB_2632707) in PBS at 4°C. Plates were washed three times with PBS immediately prior to addition of T cells. Clone 4 CTLs or TILs were resuspended in complete medium at a concentration of 1.5×10^6^ cells/mL and incubated at 37°C 100μL of CTLs or TILs was added to the well, along with 100μL of complete medium. Cells were centrifuged in the plate for 30 seconds at 250xg to ensure uniform contact with the anti-CD3 coated plastic. The plate was then placed immediately at 37°C for either 1, 2, or 5 minutes. After incubation, medium was aspirated without disturbing the CTL monolayer and cells were lysed with 100μL ice-cold RIPA buffer. Phos-tag reagent (Wako) was added to 15% SDS-PAGE gels, as per manufacturers protocols. Standard ECL protocols were used for immunodetection. Antibodies used were anti-cofilin (D3F9, Cell Signaling, RRID: AB_10622000), anti-GAPDH (14C10, Cell Signaling, RRID: AB_10693448) and anti-IgG-HRP (Cell Signaling).

### Analysis of surface receptor staining by flow cytometry

Cells to be fixed were stained with Zombie live/dead fixable exclusion dyes (biolegend). Cells were incubated with Fc receptor blocking antibody (anti-CD16/CD32, clone 93, eBioscience) prior to staining. Surface staining was conducted in PBS + 0.5% FBS. Cells were fixed with 4% PFA using standard protocols. Antibodies used for cell surface staining included: anti-CD279/PD-1 BV785 (29F.1A12, Biolegend, RRID: AB_2563680); anti-CD274/PD-L1 BV421 (MIH5, BD Bioscience, cat no 564714); anti-CD90.1/Thy1.1 PerCP Cy5.5 (OX-7, Biolegend, RRID: AB_961437); anti-Vβ8.1/Vβ8.2 TCR FITC (KJ16-133, eBioscience, RRID: AB_465261).

### In vivo killing assay

BALB/c mice were injected with 4×10^6^ primed Clone 4 CTLs, or PBS alone, by tail vein injection. On the second day after adoptive transfer, BALB/c splenocytes were pulsed with or without K^d^HA peptide at a final concentration of 2μg/mL for 1 hour, then washed and resuspended in PBS. K^d^HA pulsed splenocytes were then stained with 5μM CellTrace violet (CTV) (high), whilst un-pulsed splenocytes were stained with 0.5μM CTV (low). Splenocytes were washed and resuspended in PBS. Immediately prior to injection, high and low stained splenocytes were mixed in equal numbers, and 5×10^6^ cells were injected into BALB/c mice which had previously received Clone 4 CTLs or PBS. After 24 hours, spleens were harvested and red blood cells removed by ACK lysis. Splenocytes were analyzed by flow cytometry and gated on live/dead dye exclusion and CTV staining to compare numbers of CTV^high^ K^d^HA pulsed splenocytes and CTV^low^ control splenocytes.

## Acknowledgements

We acknowledge the University of Bristol FACS and Wolfson BioImaging facilities for providing equipment and technical support.

## Author Contributions

Conceptualization, Methodology, Formal Analysis and Writing – Review and Editing, RA, DJM and CW; Investigation, RA, GE, GT, DJM and CW; Writing – Original Draft and Visualization, RA and CW; Supervision and Funding Acquisition, DJM and CW

## Supplementary Materials

**Fig. S1. Clone 4 CTLs efficiently kill RencaHA cells in vivo**

(A) In vivo killing assay. BALB/c mice received 4×10^6^ Clone 4 CTLs via i.v. injection. After 48hrs, mice received 2.5×10^6^ of HA pulsed (CTV high) and 2.5×10^6^ of un-pulsed (CTV low) syngeneic splenocytes. Data shows the relative ratio of high vs low CTV cells isolated from spleens after 24 hours. Splenocytes gated on live/dead dye exclusion and CTV labelling.

**Fig. S2. Diminished MTOC polarization of Clone 4 TILs towards the T cell:APC interface**

(A) Tubulin-GFP transduced Clone 4 CTLs and TILs coupled to HA pulsed Renca^+HA^ cells were analysed overtime to determine the average position of the MTOC within the cell before and after tight cell coupling, time 0s. T cells were divided into equal thirds, detailed in inset schematic, and this was used to classify MTOC position at each time. Position was classed as 1.5 or 2.5 if the MTOC fell on a division line. Mean ± SEM, 51 CTL couples, 55 TIL couples, from ≥2 separate experiments. * p < 0.05,

** p < 0.01; p values calculated using proportions z-test.

**Fig. S3. Impaired formation of the Clone 4 TIL F-actin peripheral ring**

(A) Representative interaction over time of F-tractin-GFP transduced Clone 4 CTLs with Renca^+HA^ cells, following CTL adoptive transfer into RencaHA tumour bearing mice and re-isolation from the spleen after four days. Shown are matching DIC and maximum GFP projection panels with a false colour GFP intensity scale (blue to red). Scale bar shows 5μm.

(B, C) Percentage over time of F-tractin-GFP transduced Clone 4 CTLs, TILs and transferred CTLs re-isolated from the spleen, displaying peripheral accumulation of F-actin-GFP during interaction with HA peptide pulsed Renca^+HA^ cells. Tight cell coupling: time 0s. Mean ± SEM, 60 CTLs, 51 TILs, 30 transferred CTLs re-isolated from spleen, from ≥2 separate experiments. Data for CTLs and TILs also shown in Fig. 3B. P values calculated by proportions z test (C).

(D-G) Percentage over time of F-tractin-GFP transduced Clone 4 T cells displaying accumulation of F-tractin-GFP into one of six interface accumulation patterns, previously described (22, 30), during interaction with HA peptide pulsed Renca^+HA^ cells. Graphs show data for CTLs (D), TILs (F), CTLs treated with 40nM jasplakinolide for 30 minutes prior to and during coupling (G), and CTLs isolated from the spleen of a tumour bearing mouse four days following adoptive transfer (E). Tight cell coupling: time 0s. Mean ± SEM for a minimum of 50 cell couples (D,F), 40 cell couples (G) and 30 cell couples (E), from ≥2 separate experiments per condition.

(H) The mean fluorescence intensity of F-tractin-GFP was measured across the entire T cell and at the immune synapse, defined as quarter of the T cell length adjacent to the immune synapse, for Clone 4 CTLs and TILs interacting with HA peptide pulsed Renca^+HA^ cells. The ratio of F-tractin-GFP fluorescence intensity between the immune synapse and entire cell was calculated. Mean ± SEM, 33 CTLs and 28 TILs analysed from ≥2 separate experiments. p values calculated using Student’s t test.

* p < 0.05, ** p < 0.01

**Fig. S4. Cofilin and Chronophin, but not Arp2/3 and Coronin 1A, displays more sustained interface accumulation in TILs**

(A, B) Percentage over time of cofilin-GFP transduced Clone 4 CTLs (A) and TILs (B) displaying accumulation of cofilin-GFP into one of six interface accumulation patterns, previously described (22, 30), during interaction with HA peptide pulsed Renca^+HA^ cells. Tight cell coupling: time 0s. Mean ± SEM of 90 CTLs and 58 TILs from ≥2 separate experiments.

(C, D) Percentage over time of Arp3-GFP transduced Clone 4 CTLs (C) and TILs (D) displaying accumulation of Arp3-GFP into one of six interface accumulation patterns, previously described (22, 30), during interaction with HA peptide pulsed Renca^+HA^ cells. Tight cell coupling: time 0s. Mean ± SEM of 56 CTLs and 51 TILs from ≥2 separate experiments

(E) Full version of the representative western blot from Fig. 4.

(F, G) Representative interaction over time of Chronophin-GFP transduced Clone 4 CTLs (F) and TILs (G) with HA peptide pulsed Renca^+HA^ cells. Shown are matching DIC and maximum GFP projection panels with a false colour GFP intensity scale (blue to red). Scale bar shows 5μm. Complete CTL interaction in Videos S8.

(H, I) Percentage over time of Chronophin-GFP transduced Clone 4 CTLs (H) and TILs (I) displaying accumulation of Chronophin-GFP into one of six interface accumulation patterns, previously described (22, 30), during interaction with HA peptide pulsed Renca^+HA^ cells. Tight cell coupling: time 0s. Mean ± SEM of 58 CTLs and 41 TILs from ≥2 separate experiments

(J, K) Representative interaction over time of Coronin 1A-GFP transduced Clone 4 CTLs (J) and TILs (K) with HA peptide pulsed Renca^+HA^ cells. Shown are matching DIC and maximum GFP projection panels with a false colour GFP intensity scale (blue to red). Scale bar shows 5μm. Complete CTL interaction in Videos S9.

(L, M) Percentage over time of Coronin 1A-GFP transduced Clone 4 CTLs (L) and TILs (M) displaying accumulation of Coronin 1A-GFP into one of six interface accumulation patterns, previously described (22, 30), during interaction with HA peptide pulsed Renca^+HA^ cells. Tight cell coupling: time 0s. Mean ± SEM of 52 CTLs and 60 TILs from ≥2 separate experiments

**Fig. S5. SHP-1 displays more sustained interface accumulation in TILs**

(A-C) Percentage over time of Clone 4 CTLs, transduced with LAT-GFP (A), TCRζ-GFP (B) and PIP2-GFP (C), displaying accumulation of GFP-sensor into one of six interface accumulation patterns, previously described (22, 30), during interaction with HA peptide pulsed Renca^+HA^ cells. Tight cell coupling: time 0s. Mean ± SEM. (A) 55 cell couples, (B) 29 cell couples, (C) 32 cell couples, from ≥2 separate experiments per condition. Representative videos are videos S12-14.

(D) Representative interaction over time of SHP-1-GFP transduced Clone 4 TILs with HA peptide pulsed Renca^+HA^ cells. Shown are matching DIC and maximum GFP projection panels with a false colour GFP intensity scale (blue to red). Scale bar shows 5μm. Complete interaction in Video S15. (E, F) Percentage over time of SHP-1-GFP transduced Clone 4 CTLs (E) and TILs (F) displaying accumulation of SHP-1-GFP into one of six interface accumulation patterns, previously described (22, 30), during interaction with HA peptide pulsed Renca^+HA^ cells. Tight cell coupling: time 0s. Mean ± SEM of 53 CTLs and 54 TILs from ≥2 separate experiments. P values calculated by proportions z-test,

* p < 0.05, *** p < 0.001. Blue asterisks represent p values comparing CTL and TIL peripheral accumulation, black values represent comparison of overall accumulation.

**Fig. S6. Loss of PD-1 signalling in vivo, but not ex vivo, improves TIL killing ability**

(A) Representative histograms showing intensity of PD-1 expression on Clone 4 CTLs (naïve and primed) and Clone 4 TILs from RencaHA tumours, assessed by flow cytometry.

(B) CRISPR-Cas9 genome editing was used to knockout PD-L1 in RencaHA cells. Scatter plots showing expression of PD-L1 in RencaHA cells and loss of expression in RencaHA-PD-L1^-/-^ cells. PD-L1 FMO cells used for gating. Cells also gated on DRAQ7 live/dead dye exclusion.

(C) Volume of individual tumours formed by RencaHA or RencaHA PD-L1^-/-^ cells following s.c. inoculation of 1×10^6^ cells. Data compiled from two separate experiments, n=14 RencaHA and n=10 RencaHA PD-L1^-/-^.

(D) Kaplan-Meier survival curve for RencaHA and RencaHA PD-L1^-/-^ tumour bearing mice (C). P value calculated using Mantel-Cox test.

(E) Scatter plots showing PD-L1-GFP expression in RencaWT cell lines and RencaWT cells transduced with PD-L1-GFP.

**Fig. S7. Loss of PD-1 signalling in vivo significantly rescues normal F-actin and cofilin regulation**

(A, B) Clone 4 TILs were isolated from RencaHA tumour bearing mice treated with anti-PD-1 mAb, as in Fig. 5B, and from RencaHA-PD-L1^-/-^ tumour bearing mice. The percentage of cells with off-synapse lamellae (A) and the time of the first off-synapse lamella (B) during interaction with HA peptide pulsed Renca^+HA^ cells is compared to Clone 4 CTLs and TILs from RencaHA tumour bearing mice (data shown previously, fig. 2F,G). Mean ± SEM, 132 CTL, 137 TIL, 54 TIL from anti-PD-1 treated mice, 65 TIL from RencaHA-PD-L1^-/-^ tumour bearing mice. Data from ≥3 separate experiments per condition. P values calculated using one-way ANOVA.

(C, D) Percentage over time of F-tractin-GFP transduced Clone 4 TILs from anti-PD-1 mAb treated tumour bearing mice (C), and Clone 4 TILs from RencaHA-PD-L1^-/-^ tumour bearing mice (D), displaying accumulation of F-tractin-GFP into one of six interface accumulation patterns, previously described (22, 30), during interaction with HA peptide pulsed Renca^+HA^ cells. Tight cell coupling: time 0s. Mean ± SEM, 42 cells (C), 43 cells (D). Data from ≥2 separate experiments per condition.

(E, F) Percentage over time of cofilin-GFP transduced Clone 4 TILs from anti-PD-1 mAb treated tumour bearing mice (C), and Clone 4 TILs from RencaHA-PD-L1^-/-^ tumour bearing mice (D), displaying accumulation of cofilin-GFP into one of six interface accumulation patterns, previously described (22, 30), during interaction with HA peptide pulsed Renca^+HA^ cells. Tight cell coupling: time 0s. Mean ± SEM, 32 cells (E), 55 cells (F). Data from ≥2 separate experiments per condition.

(G) The mean fluorescence intensity of F-tractin-GFP was measured within the entire T cell and at the immune synapse, defined as quarter of the T cell length adjacent to the immune synapse, for Clone 4 CTLs, TILs, TILs from RencaHA tumour bearing mice treated with anti-PD-1 mAb, as in Fig. 5B, and TILs from RencaHA-PD-L1^-/-^ bearing mice, during interaction with HA peptide pulsed Renca^+HA^ cells. The ratio of f-tractin-GFP fluorescence intensity between the immune synapse and entire cell was calculated. Tight cell coupling: time 0s. Mean ± SEM, 33 CTLs, 28 TILs, 30 TILs from anti-PD-1 treated mice, 32 TILs from RencaHA-PD-L1^-/-^ bearing mice. Data from ≥2 separate experiments. P values calculated using Student’s t test with no significant difference.

** p < 0.01, *** p < 0.001, **** p < 0.0001

**Fig. S8. Acute in vitro blockade of PD-1 does not restore cytoskeletal polarization**

(A, B) Clone 4 CTLs and TILs, with and without acute in vitro anti-PD-1 treatment, coupled to HA peptide pulsed Renca^+HA^ cells, were analysed over time to determine the percentage of T cells with off-synapse lamellae (A) and the time of the first off-synapse lamella (B). Data shows mean ± SEM. Data for CTLs and TILs without anti-PD-1 treatment was previously shown in Fig. 2F, 2G.

(C, D) Percentage over time of F-tractin-GFP transduced Clone 4 CTLs (D) and TILs (D), treated for 1 hour in vitro (acute) with anti-PD-1 mAb, displaying accumulation of F-tractin-GFP into one of six interface accumulation patterns, previously described (22, 30), during interaction with HA peptide pulsed Renca^+HA^ cells. Tight cell coupling: time 0s. Mean ± SEM. (C) 41 cell couples, (D) 28 cell couples, from ≥2 separate experiments per condition.

* p < 0.05, **** p < 0.0001.

